# Enhanced food-related responses in the ventral medial prefrontal cortex in orexin-deficient patients

**DOI:** 10.1101/191544

**Authors:** Ruth Janke van Holst, Lieneke K. Janssen, Petra van Mierlo, Gert Jan Lammers, Roshan Cools, Sebastiaan Overeem, Esther Aarts

## Abstract

**Background:** Narcolepsy Type 1 is a chronic sleep disorder caused by a deficiency of orexin (hypocretin). In addition to sleep regulation, orexin is important for motivated control processes. Weight gain and obesity are common in narcolepsy. However, the neurocognitive processes associated with food-related control and overeating in orexin-deficient patients are unknown. We explored the neural correlates of general and food-related attentional control in narcolepsy patients (n=23) and healthy BMI-matched controls (n=20). In secondary analyses, we included patients with idiopathic hypersomnia (n=15) to assess sleepiness-related influences.

**Methods:** We measured attentional bias to food words with a Food Stroop task and general executive control with a Classic Stroop task during fMRI. Moreover, with correlational analyses, we assessed the relative contribution of the neural findings on the Food Stroop and Classic Stroop tasks to spontaneous snack intake.

**Results:** Relative to healthy controls, narcolepsy patients showed enhanced ventral medial prefrontal cortex responses and connectivity with motor cortex during the Food Stroop task, but attenuated dorsal medial prefrontal cortex responses during the Classic Stroop task. The ventral medial prefrontal cortex responses on the Food Stroop task, not the dorsal medial prefrontal cortex responses on the Classic Stroop task, were a significant predictor of snack intake. Comparing the narcolepsy patients with idiopathic hypersomnia patients revealed similar results.

**Conclusions:** These findings demonstrate that orexin deficiency is associated with decreased dorsal medial prefrontal cortex responses during general executive control and enhanced ventral medial prefrontal cortex responses during food-driven attention, with the latter predicting increases in food intake.

**Statement of Significance:** Patients with orexin (hypocretin) deficient narcolepsy type-1 often suffer from obesity as well as increased food craving, in addition to the sleep symptoms. However, whether and how orexin deficiency relates to neural differences in food-directed attention is unclear. We employed a Food Stroop task during fMRI and provide experimental evidence that the ventral medial prefrontal cortex responds more strongly to food words in narcolepsy patients than in controls. The hypothesis that this mechanism contributes to weight problems in narcolepsy is strengthened by the observation that ventral medial prefrontal cortex responses during the Food Stroop task were predictive of snack intake. These mechanistic data might thus advance the development of treatment targets for obesity in narcolepsy.

## Introduction

Narcolepsy Type 1 (NT1) is a disabling sleep disorder, primarily characterized by excessive daytime sleepiness and emotionally triggered episodes of muscle weakness called cataplexy. The disorder is caused by a loss of orexin (hypocretin)–producing neurons located in the lateral hypothalamus. Orexin mediates behavior under situations of high motivational relevance, through excitatory influences on the histaminergic, monoaminergic, and cholinergic system ^1^. Interestingly, the incidence of obesity is twice as high in narcolepsy compared with the normal population ^2–4^. We recently showed that food-specific satiety had reduced effects on food choices and caloric intake in narcolepsy patients, suggesting an important functional role for orexin in human food-related control of behaviuor^5^. However, the neurocognitive processes associated with food-related control and overeating in orexin-deficient patients are unknown.

Enhanced attention towards food over non-food information (i.e. attentional bias) has been proposed to contribute to the development and/or maintenance of obesity (e.g. for a review see^6^). Functional MRI studies revealed that food cues relative to neutral cues can elicit enhanced activation of the reward regions in the mesolimbic dopamine pathway in overweight relative to healthy weight individuals ^7–9^, including the ventral medial prefrontal cortex (vmPFC), striatum, insula and amygdala, which might drive excessive attention towards food cues. Detecting food rapidly and maintaining attention on food could increase the likelihood of overeating and, in the long term, obesity ^10–12^. In addition, loss of executive control during food-related distraction has been related to obesity ^13^. Although obesity is a common symptom in narcolepsy and orexin neurons interact with the mesolimbic dopamine system ^14–16^, it is unclear whether narcolepsy patients show abnormal attentional bias toward food cues, and what neurocognitive mechanism would underlie this effect.

To investigate the effect of orexin deficiency on attentional bias for food, we used a Food Stroop task (i.e. measuring reaction times toward food words and neutral words) ^13,17^ during fMRI in narcolepsy patients compared with healthy BMI-matched controls. Since spontaneous snack intake was already shown to be increased in narcolepsy versus controls in a largely overlapping sample ^5^, we investigated whether brain responses on the Food Stroop would relate to this snack intake. Additionally, we applied a Classic Stroop task (i.e. measuring response conflict) to assess general executive control abilities and evaluated the relative contribution of the neural findings on the Classic Stroop and Food Stroop tasks to spontaneous snack intake. In secondary analyses, we also compared narcolepsy patients with a control group of patients with idiopathic hypersomnia, without orexin deficiency, to verify that our findings in narcolepsy patients were not attributable to possible decreased alertness and medication-withdrawal (patients were at least 1 week off medication).

## Methods and Materials

### Participants

Fifty-eight right-handed participants were included in the experiment (20 healthy controls, 23 narcolepsy type 1 (NT1) patients and 15 idiopathic hypersomnia (IH) patients). Patients were recruited from the outpatient clinics of Sleep Medicine Center Kempenhaeghe (Heeze, the Netherlands), Sleep-wake Center SEIN ‘(Heemstede, the Netherlands) and through advertisement by the Dutch narcolepsy patients’ organization. Healthy control participants were recruited via poster and word-of-mouth advertisements in Nijmegen and surrounding areas. Healthy controls were matched to the NT1 patients in terms of average age, gender, BMI and level of education. Recruitment of IH patients was more difficult because of the rareness of the disorder ^18^ and therefore this resulted in a smaller group relative to the NT1 and healthy control group.

Inclusion criteria were age 18-60 years old, BMI 20-35 and right-handedness. Exclusion criteria were diabetes mellitus, (a history of) clinically significant hepatic, cardiac, renal, cerebrovascular, endocrine, metabolic or pulmonary disease, uncontrolled hypertension, (a history of) clinically significant neurological or psychiatric disorders and current psychological treatment other than for narcolepsy or idiopathic hypersomnia, deafness, blindness, or sensory-motor handicaps, history of taste or smell impairments, drug, alcohol or gamble addiction in the past 6 months, inadequate command of Dutch language, current strict dieting (i.e. calorie-restricted diet and/or in treatment with dietician), or food allergy to one of the ingredients used in the experiment.

All patients were diagnosed according to the International Classification of Sleep Disorders – Third Edition (ICSD-3). All had clear-cut cataplexy as well as a low mean sleep latency (< 8 minutes) measured with the Multiple Sleep Latency Test (MSLT) and at least 2 sleep onset REM periods (SOREMPs) during MLST naps and the previous night’s diagnostic sleep study. In 13 patients, orexin cerebrospinal fluid levels were known and shown to be equal or lower than 110 pg/ml.

Patients with idiopathic hypersomnia all had clear excessive daytime sleepiness, a mean sleep latency at the MSLT of 8 minutes or less, and the symptoms were not explained by another sleep disorder.

All participants were recruited on a voluntary basis and gave written informed consent before the start of the study. The study was approved by the Ethical Committee of the Radboud university medical center (CMO Arnhem-Nijmegen) and reported in the acknowledged Dutch Trial register (www.trialregister.nl: TC=4508).

### Food Stroop task and Classic Stroop task

Subjects were instructed in both tasks before going into the scanner and were further familiarized with the task by practicing the color-button contingency and performing 10 practice trials with feedback (correct/incorrect) in the scanner. For task details see Fig. 1A. In short, subjects had to indicate the color of the word presented on the screen pressing the button reflecting that color as fast and accurately as possible. In the Food Stroop task, subjects were presented with food words and neutral words, whereas in the Classic Stroop task, subjects were presented with congruent color words (e.g. the word “GREEN” printed in green) or incongruent color words (e.g. the word “GREEN” printed in red). The tasks were programmed in Presentation software (Neurobehavioral Systems Inc. https://www.neurobs.com). All task stimuli were presented with a digital projector on a screen at the back end of the MRI scanner bore, which was visible via a mirror mounted on the head coil. Responses were made using an MRI-compatible button box. Twenty generally high-calorie, palatable food words were selected from word lists reported in previous studies ^17,19^. Food words were matched to twenty neutral words each in terms of word length, number of syllables and frequency of use according to the SUBTLEX-NL norms ^20^.

**Figure 1.**
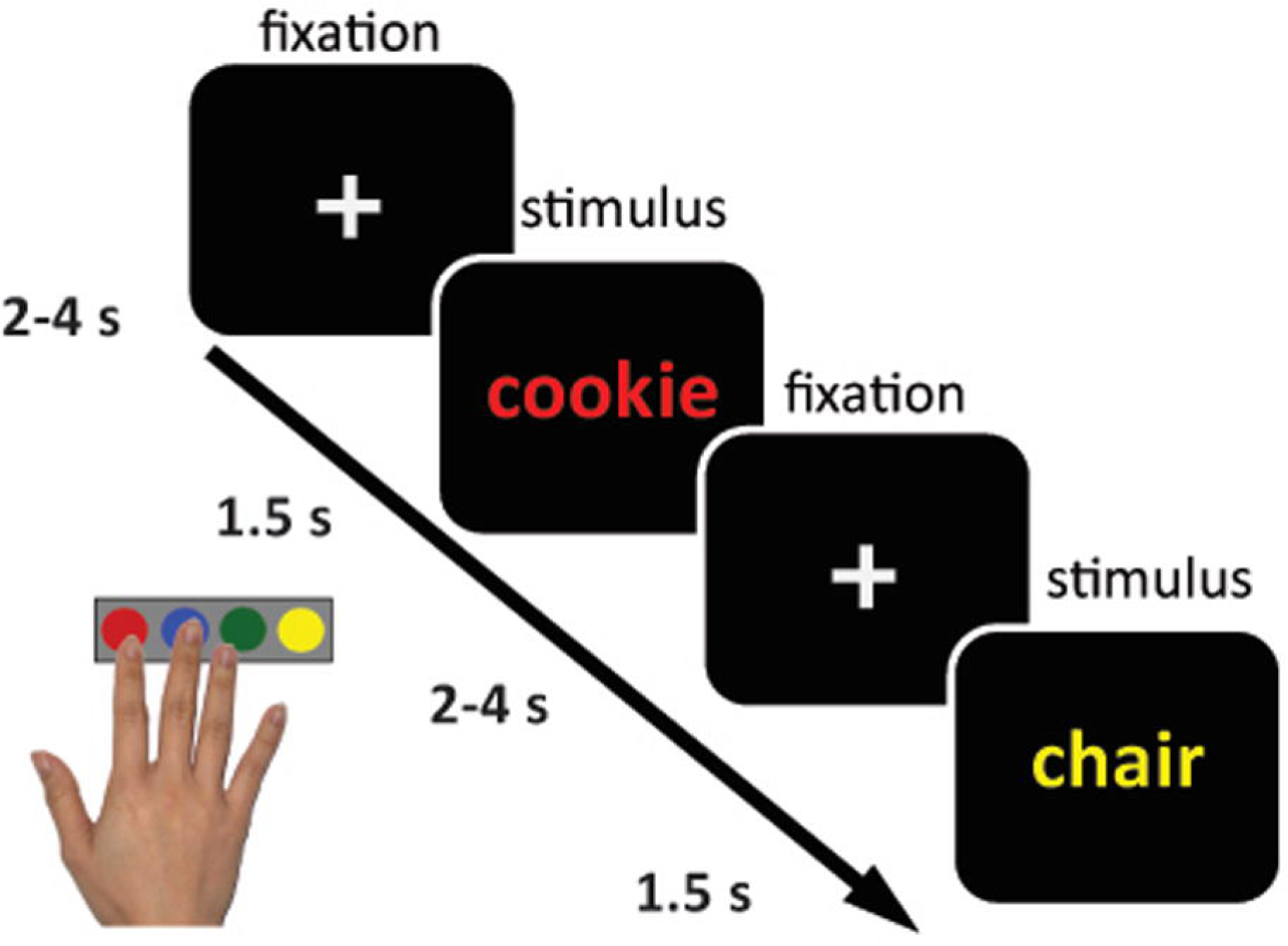
Sample trial of the Food Stroop task. On each trial, participants indicated the color of the word presented on the screen by pressing the button reflecting that color. Participants were presented with food and neutral words.

The Food Stroop interference score was calculated by subtracting the response time (RT) to neutral words from the RT to food words. Hence a higher interference score indicates more distraction by food words. Similarly, the Classic Stroop interference score was calculated by subtracting the response time (RT) to congruent words from the RT to incongruent words. Thus a higher Classic Stroop interference score indicates less general executive control ability.

### Ab-libitum snack intake

After the fMRI session, participants were asked to fill out questionnaires whilst four bowls with a variety of snacks were placed in front of them (see^5^ for the results in a largely overlapping, but larger sample). The four bowls contained: crisps, raisins, wine gums and cocktail nuts. They were told that they could eat the snacks if they felt like it. Unbeknownst to participants, we calculated the amount of kilocalories (kcal) consumed by weighting the bowl before and after, and by multiplying the amount of grams consumed by the amount of kcal/gram of that particular snack.

### Study procedure

The patient groups were asked to refrain from using their medication, if any, one week prior to the test day. On the day preceding the test day, all participants had to refrain from alcohol and drug intake, and participants had to refrain from smoking on the test day itself. Furthermore, participants fasted for at least 5 hours before the test session to ensure that they were motivated by food and snacks. The test session took place between 9am and 6pm. Timing of the session was matched between groups. During the test day (3.5 hours in total) the participants completed questionnaires (e.g. Epworth Sleepiness Scale ^21^ and Pittsburgh Sleep Quality Index ^22^ and the digit span to assess working memory capacity) and performed the Food Stroop task, directly followed by the classic color-word Stroop task during the MRI session. The test day was concluded by a behavioral satiation task and questionnaires while participants had access to ad libitum snacks; results from these measures were reported in a previous study ^5^. The number of participants included in the current analyses is smaller and not completely overlapping with the previous study because some people who did complete the satiation task did not have usable scan data (NT1 n=1 and IH=1) and vice versa (healthy controls n=1, NT1 n=1, IH=1).

### Behavioral Data Analysis

The mean latencies of the correct responses to the words and the number of correct responses in the tasks were analyzed with SPSS. We excluded trials with a RT < 200 msec.

Two narcolepsy patients (scoring 0% and 5% accuracy) and two IH patients (scoring both 10% accuracy) scored <= 10% on accuracy on the incongruent trials in the Classic Stroop task, resulting in too small number of trials to include in the fMRI analyses. These patients were therefore excluded from the Classic Stroop analyses (remaining NT1 group of n = 21 and IH group of n = 13), though they were included in the Food Stroop analyses. Behavioral group analyses including these outliers indicated no qualitatively different results on the Classic Stroop task compared with excluding these outliers (data not shown).

The median response times were used to ensure that all assumptions of parametric data were met. All behavioral outcome measures were tested for and met the homogeneity of variance assumption. Repeated measurement ANOVAs were used for the two Stroop tasks separately, to test the main effect of Condition (Food Stroop: food, neutral; Classic Stroop: incongruent, congruent), Group (NT1, healthy controls), and Group * Condition interaction effects.

Secondary analyses compared NT1 patients with IH patients using repeated measurement ANOVAs for the two Stroop tasks separately, to test the main effect of Condition (Food Stroop: food, neutral; Classic Stroop: incongruent, congruent), Group (NT1, IH), and Group * Condition interaction effects.

### Functional Imaging

Whole-brain imaging was performed on a 3 Tesla Siemens MR scanner located at the Donders Centre for Cognitive Neuroimaging, Nijmegen, The Netherlands. BOLD-sensitive functional images were acquired using a gradient-echo planar multi-echo scanning sequence (TR: 2070 ms; TEs for 4 echoes: 9 ms, 19.25 ms, 29.5 ms and 39.75 ms). We used a multi-echo EPI sequence to reduce image distortion and increase BOLD sensitivity in regions which are typically affected by strong susceptibility artifacts, such as the ventral striatum and vmPFC ^23^. One volume consisted of 34 axial slices (voxel size: 3.5 x 3.5 x 3.0 mm^3^, field of view: 224 mm, flip angle: 90°). After acquisition of the functional images, a high-resolution anatomical scan (T1-weighted MP-RAGE, TR: 2300 ms, TE: 3.03 ms, 8° flip-angle, 192 sagittal slices, slice-matrix size: 256x256, voxel size: 1x1x1 mm^3^) was obtained. Total duration of MRI sessions was 45-60 minutes.

Data were pre-processed and analyzed using SPM8 (www.fil.ion.ucl.ac.uk/spm). The volumes for each echo time were realigned to correct for motion (estimation of the realignment parameters was done for the first echo and then copied to the other echoes). The four echo images were combined into a single MR volume based on 31 volumes acquired before the actual experiment started using an optimised echo weighting method. Combined functional images were slice-time corrected by realigning the time-series for each voxel temporally to acquisition of the middle slice. Structural and functional data were then co-registered and spatially normalised to a standardized stereotactic space (Montreal Neurological Institute (MNI) template). After segmentation of the structural images using a unified segmentation approach, the mean of the functional images was spatially coregistered to the bias-corrected structural images. The transformation matrix resulting from segmentation was then used to normalize the final functional images into MNI space (resampled at voxel size 2x2x2 mm). Finally, the normalised functional images were spatially smoothed using an isotropic 8 mm full-width at half-maximum Gaussian kernel.

### Functional MRI Data Analysis

Statistical analyses were performed according a general linear model (GLM) as implemented in SPM8. At the first level, subject-specific data were analyzed using a fixed effects model which contained 2 regressors of interest with the correct trials on food trials and those on neutral trials of the Food Stroop task and 2 regressors with the correct trials on incongruent trials and those on congruent trials of the Classic Stroop task. All onsets were modeled using a stick function and convolved with the canonical hemodynamic response function. We also included regressors of non-interest: one for incorrect trials, one for missed trials, as well as six movement parameters - resulting from the realignment procedure - and their six time derivatives to account for head movement, and finally the average ‘out of brain’ signal, derived from the segmented anatomical scan. High pass filtering (128 seconds) was applied to the time series of the functional images to remove low-frequency drifts and correction for serial correlations was done using an autoregressive AR(1) model.

At the second level, we investigated whole-brain main effect of the tasks and group effects in a random effects analysis. Group differences in brain responses on the Food Stroop (food – neutral) and on the Classic Stroop (incongruent – congruent) contrast were tested with an independent two-sample t-test; the main effects of the tasks were tested with a one-sample t-test. In all second level analyses, we added as covariate of non-interest a summary motion score for every subject, which was calculated as the sum of the root-mean-square value of the subject’s frame wise-displacement parameters (x, y, z in mm & pitch, roll, and yaw in degrees ^24^. We tested for correlations between whole-brain responses to food vs neutral words and spontaneous snack intake, as well as BMI, across healthy controls and NT1 patients. Additionally, we assessed the relative contribution of the neural findings (by extracting beta’s from the relevant clusters) on the Food Stroop and Classic Stroop tasks to spontaneous snack intake across healthy control and NT1 groups by using them as predictors in a multiple regression model in SPSS using the forced entry (or Enter as it is known in SPSS and using a p<0.05 to report significant results) method. For the fMRI analyses we used an FWE-corrected cluster level threshold p < 0.05 (intensity threshold, uncorrected p<0.001).

The secondary comparisons with our extra control group, i.e. the IH patients, are described under the heading “Control comparisons” in the Results section.

### Generalized Psycho-Physiological Interaction (gPPI) Analysis

To test functional connectivity differences between groups during color-naming of food versus neutral words, we conducted a generalized psychophysiological interaction analysis ^25^. As a seed for the gPPI analyses we used the one cluster that was significantly different between the healthy controls and narcolepsy patients (see Results) during the Food Stroop task (i.e. the right ventral medial prefrontal cortex). See Figure 4a for details and the resulting seed. Because we modeled the main effect of task in the PPI analysis, the PPI will only detect functional connectivity effects over and above (orthogonal to) the main effect of task, thus there is no concern about non-independence or circularity in this case^26^.

**Figure 4.**
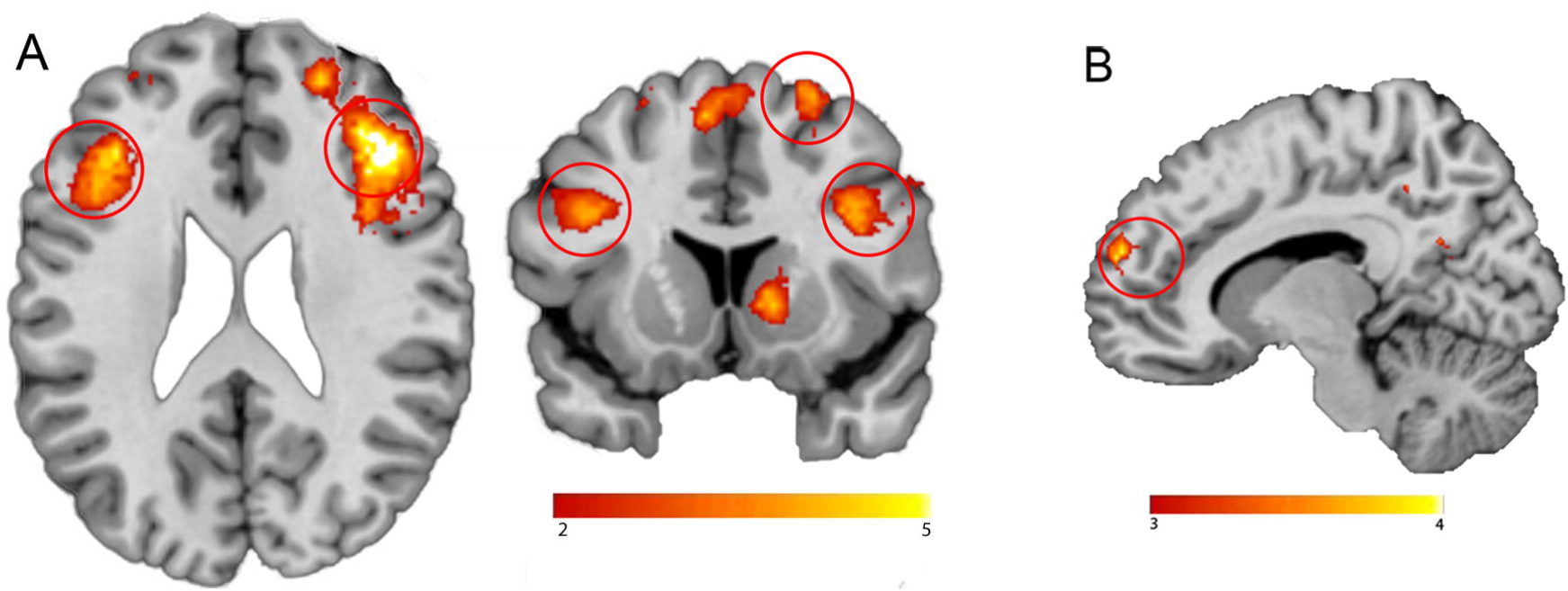
**a) The right vmPFC seed**, defined as the significant cluster from the food – neutral trials contrast indicating more activity in Narcolepsy type 1 patients relative to healthy controls (Figure 3b), combined with the corresponding Automated Anatomical Labeling (AAL) masks. **b)** Functional connectivity between the vmPFC seed and the right motor cortex was higher in Narcolepsy type 1 patients. All statistical parametric maps were overlaid onto a T1-weighted canonical image. Images are shown in neurological convention (left = left). Full brain statistical parametric maps were thresholded at p <0.001 uncorrected, encircled regions are significant clusters at pFWE<0.05. Color scale indicates T-scores ranging from 3 (red) to 4 (yellow).

We used the generalized PPI toolbox (gPPI; http://www.nitrc.org/projects/gppi; McLaren et al., 2012) in SPM8 (Statistical Parametric Mapping, Wellcome Department of Cognitive Neurology, London, UK), given that gPPI has the flexibility to accommodate multiple task conditions in the same connectivity model. To estimate the neural activity producing the physiological effect in the seed region for each subject, the BOLD signal was extracted from this region and deconvolved ^27^. This was included in the model as a physiological regressor, as were the onset times for each of the task conditions (food, neutral, congruent and incongruent words) as psychological regressors, as well as the physiological regressor multiplied by the psychological regressors (convolved with the HRF), resulting in nine regressors on the first level (i.e., one physiological, four psychological, and four interaction regressors). One PPI contrast was created for each subject: food trials – neutral trials. On the second level, this PPI contrast was analyzed using independent two-sample t-tests comparing healthy controls with narcolepsy patients. We used an FWE-corrected cluster level threshold of p < 0.05 (intensity threshold uncorrected p<0.001).

The additional comparisons with our extra control group, i.e. the IH patients, are shortly described under the heading “Control comparisons” in the Results section.

## Results

### Participants

Table 1 summarizes the demographic and clinical characteristics of the participants who were included in the data analysis. Narcolepsy patients and healthy controls were well matched on gender, age, BMI and education level. Patient groups did not differ in medication type used, daytime sleepiness, and were –as expected- significantly sleepier than the healthy controls. As expected, there was significant difference between NT1 and IH patients on the quality of sleep, with narcolepsy patients reporting a lower quality of sleep. Working memory capacity, as measured with the digit span, did not differ between the groups.

### Ad libitum food intake

Narcolepsy patients spontaneously consumed significantly more calories (mean: 324.71 SD: 272.20) during the ad-libitum snack intake than healthy controls (mean: 114.29 SD: 150.46; F(1,42)=11.108 p=0.002).

### Behavioral performance on the Food Stroop task

Participants did not respond faster to food words than to neutral words (main effect of Condition: F(1,39)=1.767, p=0.192). We did not see a main effect of Group: (F(1,39)= 0.941, p=0.338), and no significant Condition * Group effect on RTs (F(1,39)= 1.354, p=0.257) (Table 2; Figure 2).

**Figure 2.**
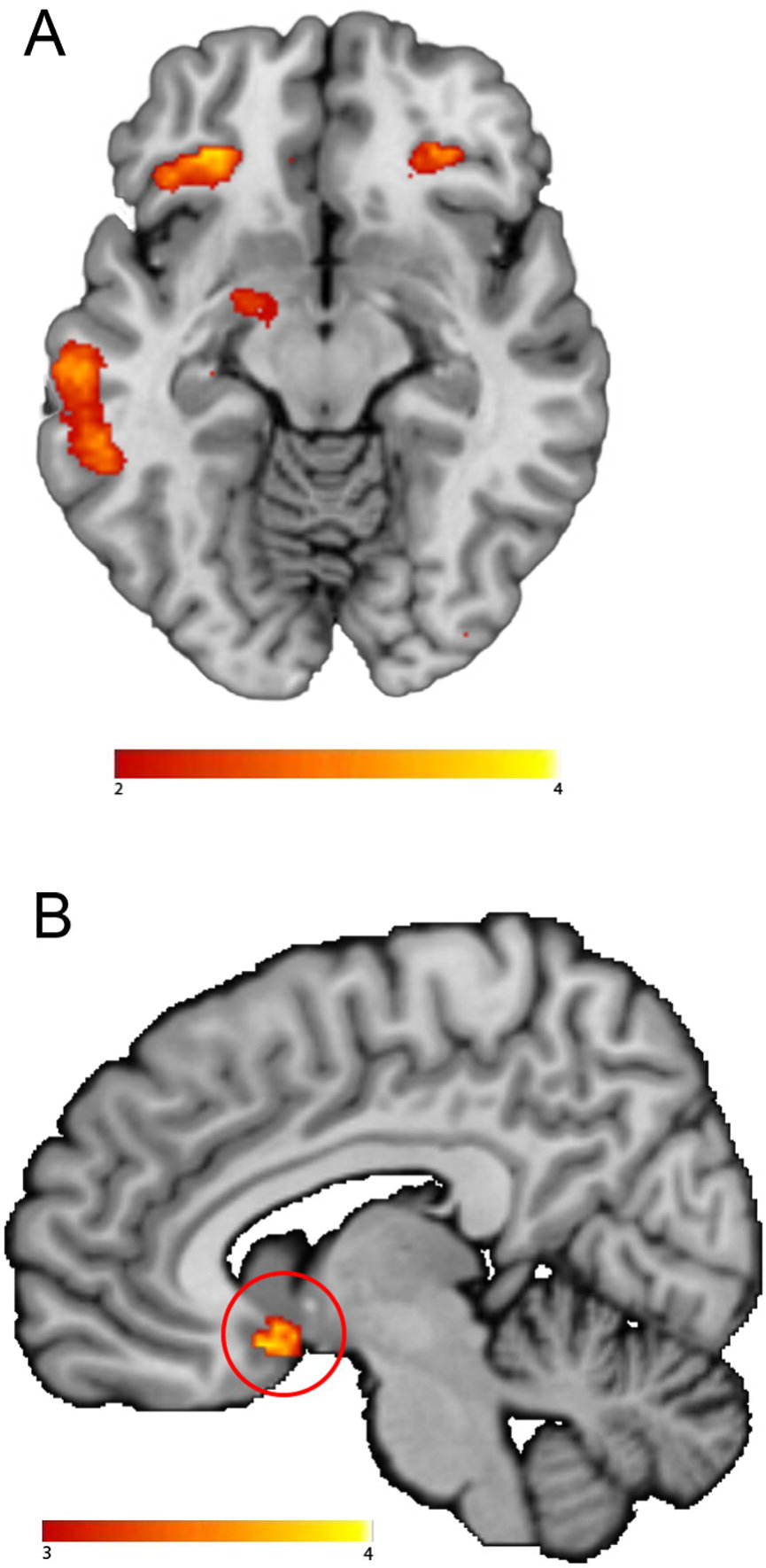
Reaction times during the Food Stroop task. Values are RT means for the difference between food and neutral words. Bars indicate standard errors of the group mean. HC= healthy controls; NT1 patients= Narcolepsy type 1 patients; IH: idiopathic hypersomnia patients. RT= reaction time; ms= milliseconds.

For Food Stroop accuracy, we observed no main effect of Condition (F(1,395)=0.954, p=0.335), no main effect of Group (F(1,39)=0.005, p=0.947), and no significant Condition * Group effect (F(1,39)=1.050, p=0.312).

### Food Stroop fMRI results

The main task effect of the contrast food words minus neutral words across groups yielded no significant brain responses when applying the pFWE<.05 cluster corrected threshold. Using an uncorrected threshold (p<.001), responses were found in the right inferior frontal cortex (Brodmann area 48; x,y,z: 42, 30, 16, t=4.32, k=45, p_cluster_uncorrected_=0.006), left inferior orbitofrontal cortex (Brodmann area 11; x,y,z: -30, 34,-14, t=4.10, k=40, p_cluster_uncorrected_ =0.009; Figure 3a) and in the left hippocampus (Brodmann area 20 x,y,z: -30, -20, -18, t=4.56, k=21, pcluster_uncorrected =0.047). Importantly, on a corrected threshold, NT1 patients displayed increased responses for food versus neutral words in a region of the reward circuitry, i.e. the ventral medial prefrontal cortex (vmPFC/Brodmann area 25; x,y,z: 6, 10,-14, t=4.45, k=87, p_cluster-FWE_=0.011; Figure 4b) compared with healthy controls. We did not observe significant correlations with BMI scores nor snack intake, within or across groups. If anything, snack intake correlated positively with a cluster in the vmPFC at p<0.001 uncorrected, but this did not survive multiple comparison correction (x,y,z: 26, 18, -16, t=4.18, k=28, p_cluster-FWE_=0.543).

**Figure 3.**
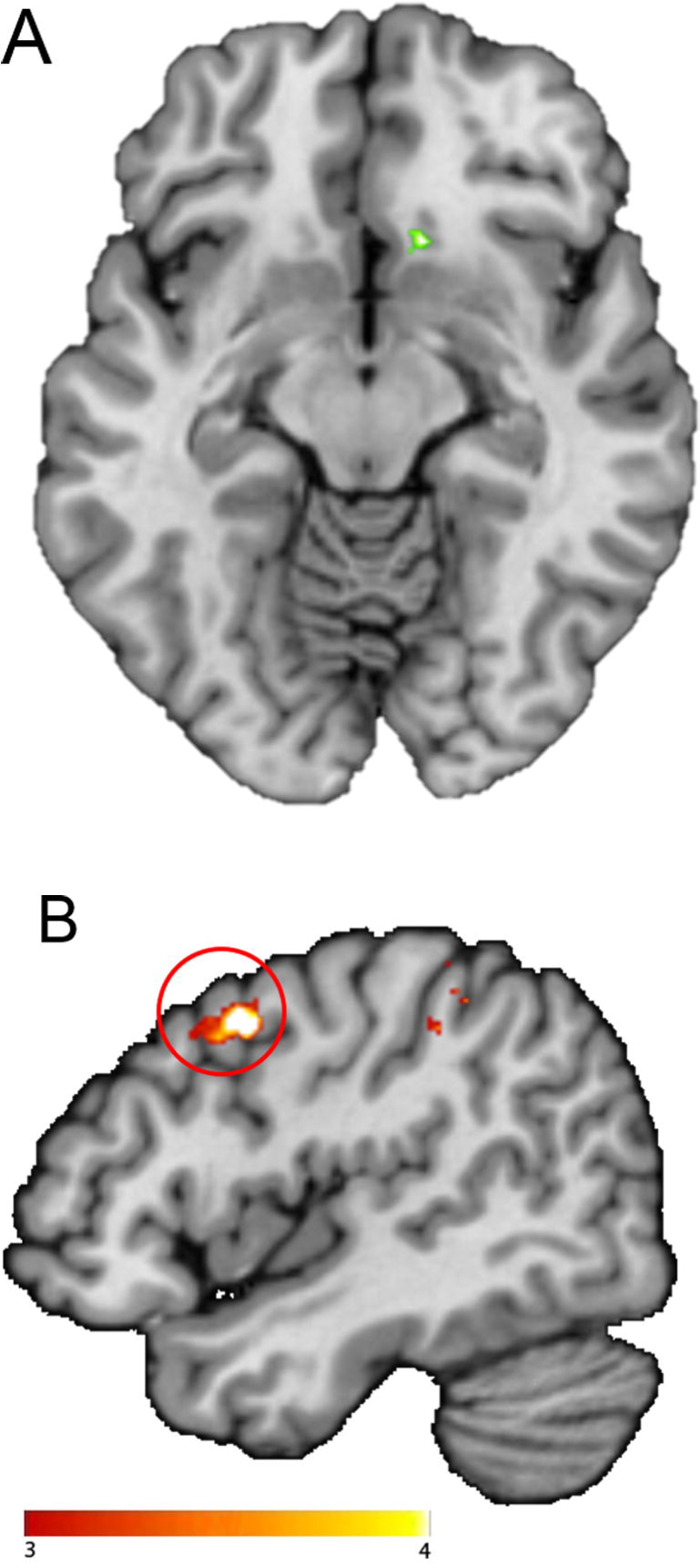
Neural Food Stroop effect. a) Main effect of the contrast of food minus matched neutral words. b) Stronger BOLD response in Narcolepsy type 1 patients versus healthy controls on the food versus neutral words contrast. All statistical parametric maps were overlaid onto a T1-weighted canonical image. Images are shown in neurological convention (left = left). Full brain statistical parametric maps were thresholded at p <0.001 uncorrected, encircled regions are significant clusters at pFWE<.05. Color scale indicates T-scores ranging from 3 (red) to 4 (yellow).

### Functional connectivity with the vmPFC seed during the Food Stroop task

We found stronger functional connectivity for NT1 patients relative to healthy controls between the vmPFC seed and the right premotor cortex during the Food Stroop task (food – neutral trials) (Brodmann area 6; x,y,z: 46, 8, 42, t=4.48, k=70, p_cluster-FWE_=0.032; Figure 4b).

### Behavioral performance on the Classic Stroop task

To check whether any observed differences in the Food Stroop task or in snack intake could be due to general executive control deficits^28^, we employed the Classic Stroop task. As expected, participants were faster on the congruent trials than on the incongruent trials (main Condition: F(1,39)=11.691, p=0.001). We did not see a main effect of Group (F(1,39)=0.533, p=0.470), and no significant Condition * Group effect on RTs (F(1,39)=0.004, p=0.949) (Table 2). Participants were also more accurate on the congruent versus the incongruent trials (main Condition: F(1,39)=4.097, p=0.05). We did not see group differences across trials (main Group: F(1,39)=0.157, p=0.694) or as a function of congruency (Condition * Group: F(1,39)=0.258, p=0.614) (Table 2).

### Classic Stroop fMRI results

The main task effect of the contrast incongruent words minus congruent words across groups resulted in significant clusters (Figure 5a) in the bilateral inferior frontal cortex (right x,y,z: 40, 26, 22, t=5.89, k=773, p_cluster-FWE_<0.001; left x,y,z: -36, 24, 20, t=5.02, k=725, p_cluster-FWE_<0.001), supplementary motor cortex (x,y,z: -6, 14, 54, t=4.87, k=201, p_cluster-FW_E<0.001), right superior frontal cortex (x,y,z: 26, 12, 60, t=4.82, k=233, p_cluster-FW_E<0.001) and left middle frontal cortex (x,y,z: -26, -10, 52, t=4.75, k=85, p_cluster-FWE_=0.014). Compared with healthy controls, NT1 patients displayed lower responses for incongruent words minus congruent words in the left dorsal medial prefrontal cortex (dmPFC) (superior frontal gyrus/ Brodmann area 32; x,y,z: -18, 38, 34, t=5.73, k=191, p_cluster-FWE_<0.001: Figure 5b).

**Figure 5.**
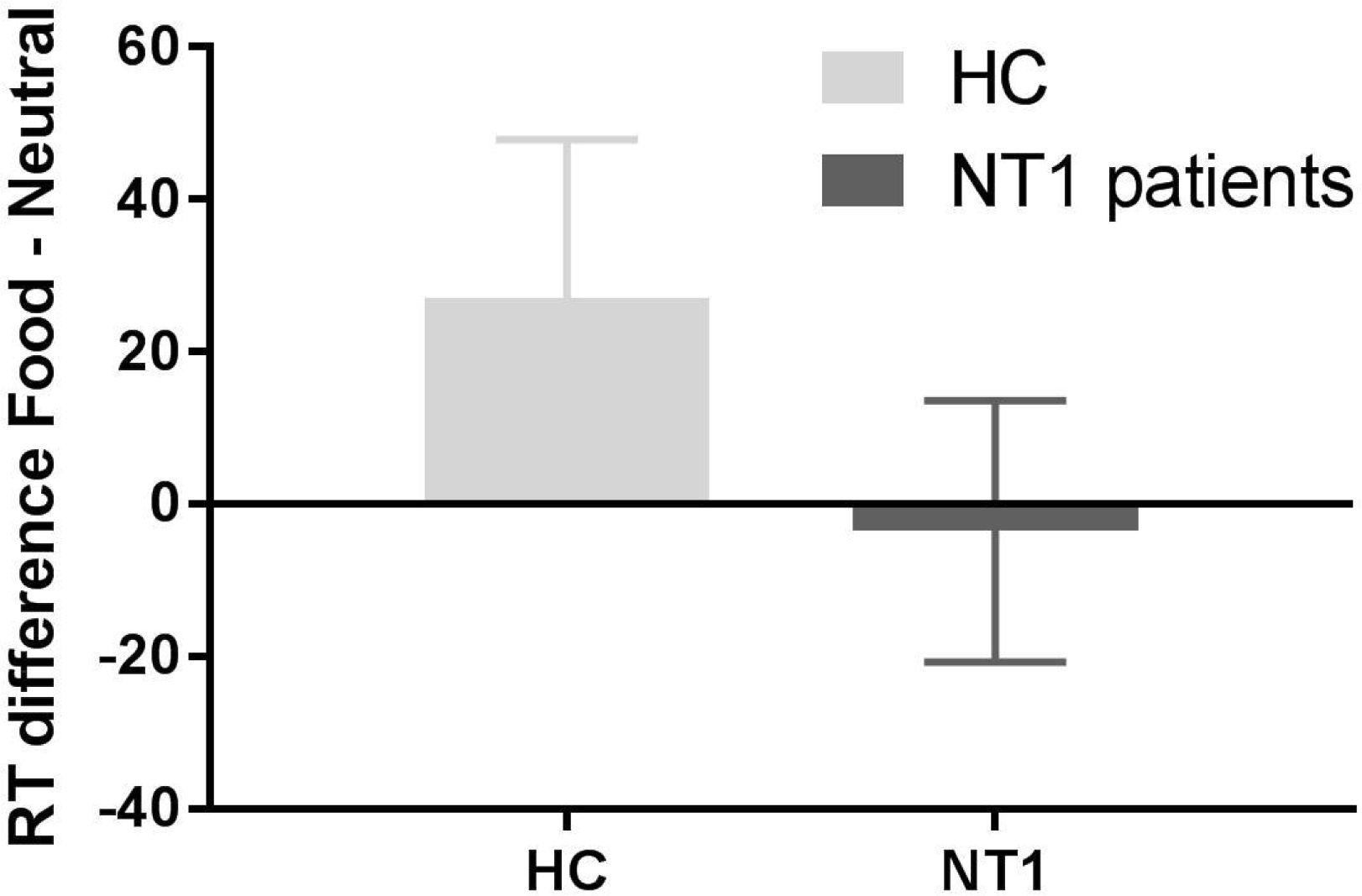
**a)** Main effect across groups on the incongruent versus congruent words contrast in the classic Stroop task. Color scale indicates T-scores ranging from 2 (red) to 5 (yellow). **b)** Stronger BOLD response in healthy controls versus Narcolepsy Type 1 patients on the incongruent versus congruent words contrast. All statistical parametric maps were overlaid onto a T1-weighted canonical image. Images are shown in neurological convention (left = left). Full brain statistical parametric maps were thresholded at p <0.001 uncorrected (for illustration purposes), encircled regions are significant clusters at pFWE<0.05. Color scale indicates T-scores ranging from 3 (red) to 4 (yellow).

### Relative contribution of neural Stroop responses in predicting food intake

To assess the relative contribution of the responses in the vmPFC responses during the Food Stroop task and in the dmPFC during the Classic Stroop task to spontaneous snack intake, we performed a multiple regression analysis across participants using the beta values extracted from the vmPFC and dmPFC clusters. A regression model was found (F(2,38) = 3.127, p=0.056), with R^2^ of .148. Only the vmPFC responses elicited on the Food Stroop task were a significant positive predictor of spontaneous snack intake (standardized Beta = .330, t=2.086, p =0.043; Figure 6), whereas the dmPFC responses on the Classic Stroop task were not significantly correlated to spontaneous snack intake (standardized Beta = -0.138, t=-0.875, p =0.388). Thus, food reward-related vmPFC responses on the Food Stroop task, which were increased in NT1 patients versus healthy controls, have a relatively larger contribution to spontaneous snack intake than the executive functioning-related dmPFC cortex responses, which were decreased in narcolepsy patients versus healthy controls. Brain responses in the vmPFC and dmPFC did not significantly predict BMI scores.

**Figure 6.**
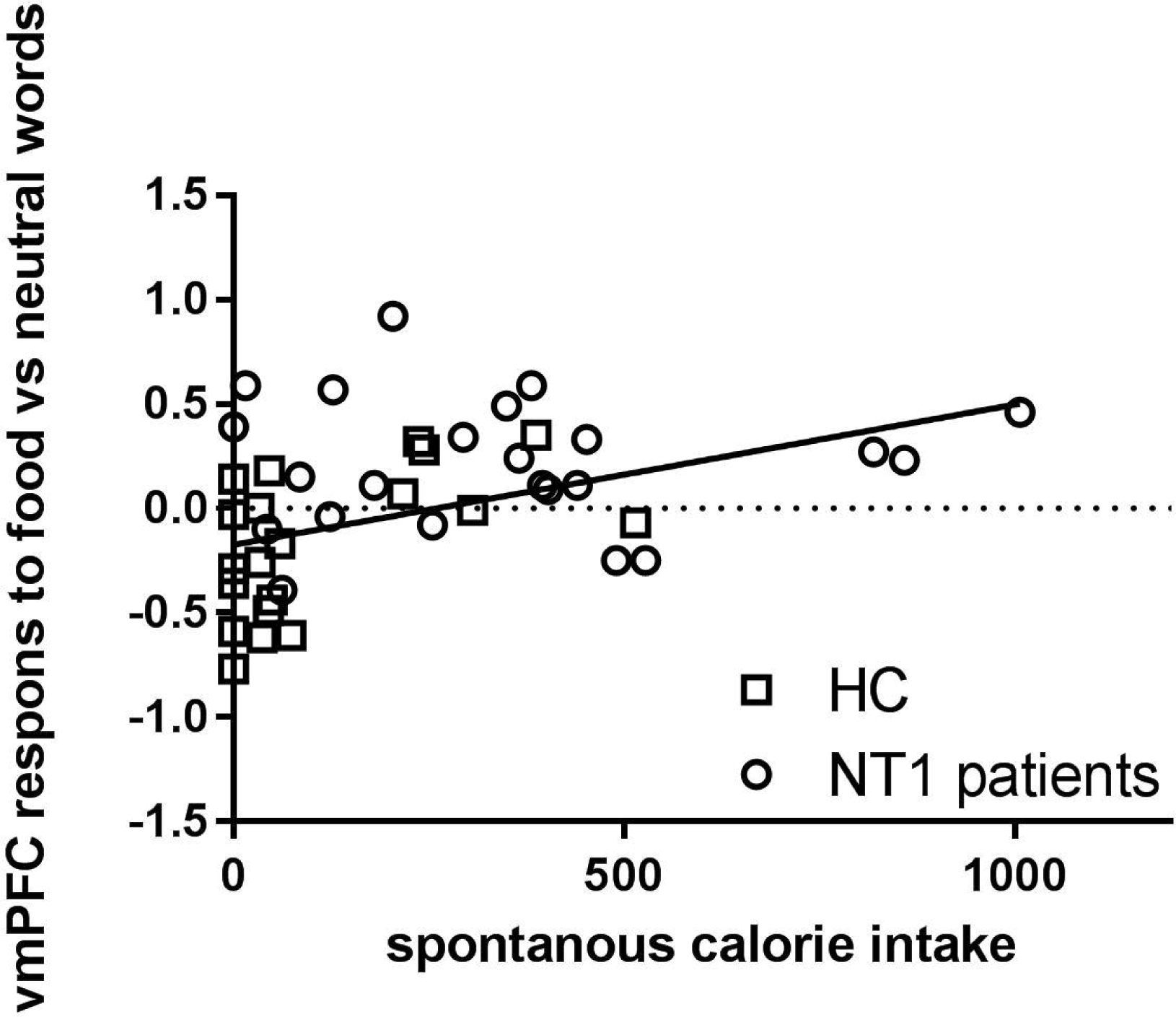
**Visual presentation of the relative contribution of the responses in the vmPFC responses during the Food Stroop task** to spontaneous snack intake in the healthy controls, Narcolepsy type 1 and idiopathic hypersomnia patients.

### Control comparisons

As in the comparison with healthy controls, NT1 patients consumed significantly more calories than IH patients (mean: 80.85 SD: 126.56; F(1,37)=12.086, p=0.001) after the task.

Similar to the comparison between healthy controls and NT1 patients, during the Food Stroop task, there were no differences in behavioural performance (Table S1) between NT1 patients and IH patients, but the vmPFC region (vmPFC/Brodmann area 25; x,y,z: 8, 14, -12, t=5.07, k=149, p_cluster-FWE_<0.001) and the left superior temporal lobe (Brodmann area 48; x,y,z: -44, -12, -08, t=5.36, k=248, p_cluster-FWE_<0.001) were more active in NT1 compared with IH patients. Moreover, we did not observe significant whole brain correlations with BMI scores nor snack intake, within or across NT1 and IH patients. In contrast to the NT1 versus HC comparisons, we found no significant between-group differences in functional connectivity with the vmPFC when comparing narcolepsy patients with IH patients.

During the Classical Stroop task there were no differences in behavioural performance nor in brain responses between NT1 patients and IH patients (Table S1).

To assess the relative contribution of the responses in the vmPFC responses during the Food Stroop task and in the dmPFC during the Classic Stroop task to spontaneous snack intake, we also performed a multiple regression analysis. A regression model was found (F(2,33) = 2.868, p=0.072), with R^2^ of .156. Only the vmPFC response elicited on the Food Stroop task revealed itself as a trending positive predictor of spontaneous snack intake (standardized Beta = .293, t=1.769, p =0.087), whereas the dmPFC response on the Classic Stroop task was not significantly correlated to spontaneous snack intake (standardized Beta = -0.244, t=-1.473, p =0.151). Brain responses in the vmPFC and dmPFC did not significantly predict BMI scores.

## Discussion

In this study we aimed to elucidate the role of orexin in neurocognitive mechanisms underlying food attentional bias by investigating orexin-deficient narcolepsy patients. Narcolepsy patients showed increased activation of the vmPFC when responding to food words relative to neutral words, compared with healthy controls as well as with IH patients (who are comparably sleepy (although having slightly higher quality of sleep) and medicated, but have normal orexin levels compered with NT1 patients). In addition, narcolepsy patients relative to healthy controls displayed higher vmPFC connectivity with the motor cortex when responding to food words relative to neutral words.

The vmPFC is part of the fronto-striatal reward circuitry and is often found to show enhanced activity when people are cued with high caloric food cues (e.g. pictures or words) versus low caloric food cues ^6,7,9,29,30^. Indeed, enhanced reactivity to food cues has been shown to predict future weight gain in healthy weight individuals ^10,11^. For example, Stice and colleagues ^12^ found that elevated vmPFC/orbitofrontal cortex responses to cues signaling impending milkshake receipt predicted future body fat gain over 3-years follow-up in healthy weight adolescents. The current finding of enhanced vmPFC responses in response to rewarding food (versus neutral) words and the enhanced functional connectivity between vmPFC and motor cortex in narcolepsy patients, suggests that narcolepsy patients have enhanced reward-driven invigoration in response to food words which could underlie their weight gain over time. This is further substantiated by our findings that the vmPFC responses to food (versus neutral) words were a positive and unique predictor (relative to dmPFC responses for incongruent versus congruent words) of spontaneous snack intake after scanning in a marginally significant regression model.

Enhanced responses to food cues have been associated with increased dopamine release in reward-related brain areas ^31,32^. Similarly, animal studies have demonstrated that, in response to salient events, orexin projections enhance dopamine firing rates in reward-related areas, including the vmPFC, nucleus accumbens and the dopaminergic ventral tegmental area (VTA) ^16,33^. Since narcolepsy is characterized with orexin deficiency, a *decrease* (via lower VTA activity) instead of an increase in activity of reward-related brain areas might have been expected. Indeed, in a study with monetary reward cues, narcolepsy patients relative to healthy controls, lacked VTA and vmPFC activity when prompted with high versus low incentive monetary cues^34^. Presently, we only observed enhanced vmPFC responses and connectivity for (food) reward-related stimuli and no accompanying diminished responses in other reward regions. How orexin deficiency in narcolepsy patients exactly relates to enhanced vmPFC activity in response to food stimuli requires further study.

On the general executive control task, e.g. Classic Stroop, narcolepsy patients displayed lower responses in dmPFC for incongruent colored words versus congruent colored words than healthy controls (but not versus IH). The dmPFC is part of the executive control network and is sensitive to the degree of response conflict ^35–37^. Behaviorally, there were no significant group differences on the Stroop interference effect, which is in line with previous cognitive studies assessing executive functioning in narcolepsy ^38,39^. Similar to narcolepsy patients, IH patients showed normal Stroop and there was no group difference in brain responses. Hence, general sleepiness in both patient groups might be related to diminished responses in these executive control/attention regions, as shown before during sleep deprivation ^40–42^. It is less likely that diminished executive control responses in dorsal frontal regions in NT1 lead to overeating, as we currently did not find a relation with snack intake.

One caveat of this study is the absence of expected Food Stroop main effects, in both behavioral and fMRI responses. A previous fMRI study used the same Food Stroop task to measure attentional bias to food words relative to neutral words in healthy controls (n=76, 85% women, BMI: 19-35) ^13^. They reported activation patterns in frontal-parietal areas (including the inferior frontal cortex, inferior orbitofrontal cortex and middle temporal cortex) and slower reaction times (i.e. indicating interference of food words) when healthy controls responded to food relative to neutral words. On a lower statistical threshold (p<0.001 uncorrected), we indeed find similar brain areas for the food vs neutral contrast as reported in Janssen et al. ^13^. The decrease in power might be due to the fact that our participants were less weight concerned than the subjects of Janssen et al^13^, who all signed up for an intervention study to change eating habits. Indeed, individuals who are preoccupied with a healthy weight also show increased behavioral food attentional bias^43,44^. Moreover, our study included both patients and healthy volunteers, with the healthy controls showing - if anything - RT interference by the food words (as in Janssen et al. ^13^), whereas the patients demonstrated - if anything - RT facilitation by food words (see Fig 2). Although these opposite behavioral effects in patients versus controls did not reach significance, they could have resulted in the absence of main task effects. Importantly, our study was able to pick up enhanced vmPFC responses in narcolepsy patients relative to healthy controls and IH patients.

One of the strengths of the current study is that we tested narcolepsy and IH patients at least 1 week off their medication, reducing the effects of, amongst others, psychostimulants in our findings. Moreover, by including a group of IH patients, we could discern sleep-disorder related issues (like excessive sleepiness and medication withdrawal) from orexin-deficiency effects. Indeed, similar behavioural and neural responses on the Food Stroop task were found when comparing narcolepsy patients with IH patients, suggesting it is unlikely that decreased alertness or medication withdrawal alone would explain our findings in narcolepsy. Our study is the first to study the neurocognitive mechanisms of food cues processing in orexin-deficient narcolepsy patients. These findings do not only point to an important role for orexin in food-related motivation in humans, but also suggest possible underlying factors of overeating in narcolepsy.

## Acknowledgment

The authors would like to thank all the patients and their families for participating in our study.

## Author Contributions

RJ.v.H. participated in the design of the experiments, performed the human studies, analyzed and interpreted most data, and wrote the manuscript.

LJ.J. participated in the design of the experiments and helped analyze the data.

P.v.M. recruited patients and helped organize the experiments.

GJ.L. recruited patients and was involved in revising the final version.

R.C. initiated the study and was involved in the experimental design, data interpretation and in revising the final version.

S.O. initiated the study and was involved in the experimental design and data interpretation. He supervised the project and was involved in revising the final version.

E.A. initiated the study, was involved in the experimental design and data interpretation. She supervised the project and was involved in revising the manuscript.

## References

1. Sakurai T. The role of orexin in motivated behaviours. Nat Rev Neurosci. 2014;15(11):719–31. doi:10.1038/nrn3837.

2. Schuld A, Hebebrand J, Geller F, Pollmächer T. Increased body-mass index in patients with narcolepsy. Lancet. 2000;355(9211):1274–5. doi:10.1016/S0140-6736(05)74704-8.

3. Dauvilliers Y, Arnulf I, Mignot E. Narcolepsy with cataplexy. Lancet. 2007;369(9560):499–511. doi:10.1016/S0140-6736(07)60237-2.

4. Kok SW, Overeem S, Visscher TLS, et al. Hypocretin deficiency in narcoleptic humans is associated with abdominal obesity. Obes Res. 2003;11(9):1147–54. doi:10.1038/oby.2003.156.

5. van Holst RJ, van der Cruijsen L, van Mierlo P, et al. Aberrant Food Choices after Satiation in Human Orexin-Deficient Narcolepsy Type 1. Sleep. 2016;39(11): 1951–1959. doi:10.5665/sleep.6222.

6. Hendrikse JJ, Cachia RL, Kothe EJ, McPhie S, Skouteris H, Hayden MJ. Attentional biases for food cues in overweight and individuals with obesity: a systematic review of the literature. Obes Rev. 2015;16(5):424–32. doi:10.1111/obr.12265.

7. Stoeckel LE, Weller RE, Cook EW, Twieg DB, Knowlton RC, Cox JE. Widespread reward-system activation in obese women in response to pictures of high-calorie foods. Neuroimage. 2008;41(2):636–47. doi:10.1016/j.neuroimage.2008.02.031.

8. Rothemund Y, Preuschhof C, Bohner G, et al. Differential activation of the dorsal striatum by high-calorie visual food stimuli in obese individuals. Neuroimage. 2007;37(2):410–21. doi:10.1016/j.neuroimage.2007.05.008.

9. Scharmüller W, Übel S, Ebner F, Schienle A. Appetite regulation during food cue exposure: a comparison of normal-weight and obese women. Neurosci Lett. 2012;518(2):106–10. doi:10.1016/j.neulet.2012.04.063.

10. Demos KE, McCaffery JM, Cournoyer SA, Wunsch CA, Wing RR, E Demos K. Greater Food-Related Stroop Interference Following Behavioral Weight Loss Intervention. J Obes Weight Loss Ther. 2013;S3(5). doi:10.4172/2165-7904.1000187.

11. Yokum S, Ng J, Stice E. Attentional bias to food images associated with elevated weight and future weight gain: an fMRI study. Obesity (Silver Spring). 2011;19(9):1775–83. doi:10.1038/oby.2011.168.

12. Stice E, Burger KS, Yokum S. Reward Region Responsivity Predicts Future Weight Gain and Moderating Effects of the TaqIA Allele. J Neurosci. 2015;35(28):10316–24. doi:10.1523/JNEUROSCI.3607-14.2015.

13. Janssen LK, Duif I, van Loon I, et al. Loss of lateral prefrontal cortex control in food-directed attention and goal-directed food choice in obesity. Neuroimage. 2017; 146:148-156. doi:10.1016/j.neuroimage.2016.11.015.

14. Muschamp JW, Hollander JA, Thompson JL, et al. Hypocretin (orexin) facilitates reward by attenuating the antireward effects of its cotransmitter dynorphin in ventral tegmental area. Proc Natl Acad Sci USA. 2014;111(16):E1648–55. doi:10.1073/pnas.1315542111.

15. Choi DL, Davis JF, Magrisso IJ, Fitzgerald ME, Lipton JW, Benoit SC. Orexin signaling in the paraventricular thalamic nucleus modulates mesolimbic dopamine and hedonic feeding in the rat. Neuroscience. 2012;210:243–8. doi:10.1016/j.neuroscience.2012.02.036.

16. Mahler S V, Moorman DE, Smith RJ, James MH, Aston-Jones G. Motivational activation: a unifying hypothesis of orexin/hypocretin function. Nat Neurosci. 2014;17(10):1298–303. doi:10.1038/nn.3810.

17. Nijs IMTT, Franken IHAA, Muris P. Food-related Stroop interference in obese and normal-weight individuals: Behavioral and electrophysiological indices. Eat Behav. 2010;11(4):258–265. doi:10.1016/j.eatbeh.2010.07.002.

18. Dauvilliers Y. Differential diagnosis in hypersomnia. Curr Neurol Neurosci Rep. 2006;6(2):156–62.

19. Phelan S, Hassenstab J, McCaffery JM, et al. Cognitive Interference From Food Cues in Weight Loss Maintainers, Normal Weight, and Obese Individuals. Obesity. 2010;19(1):69–73. doi:10.1038/oby.2010.138.

20. Keuleers E, Brysbaert M, New B. SUBTLEX-NL: a new measure for Dutch word frequency based on film subtitles. Behav Res Methods. 2010;42(3):643–50. doi:10.3758/BRM.42.3.643.

21. Johns MW. Daytime sleepiness, snoring, and obstructive sleep apnea. The Epworth Sleepiness Scale. Chest. 1993;103(1):30–6.

22. Buysse DJ, Reynolds CF, Monk TH, Berman SR, Kupfer DJ. The Pittsburgh Sleep Quality Index: a new instrument for psychiatric practice and research. Psychiatry Res. 1989;28(2):193–213.

23. Poser BA, Versluis MJ, Hoogduin JM, Norris DG. BOLD contrast sensitivity enhancement and artifact reduction with multiecho EPI: Parallel-acquired inhomogeneity-desensitized fMRI. Magn Reson Med. 2006;55(6):1227–1235. doi:10.1002/mrm.20900.

24. Jenkinson M, Bannister P, Brady M, Smith S. Improved optimization for the robust and accurate linear registration and motion correction of brain images. Neuroimage. 2002;17(2):825–41.

25. McLaren DG, Ries ML, Xu G, Johnson SC. A generalized form of context-dependent psychophysiological interactions (gPPI): A comparison to standard approaches. Neuroimage. 2012;61(4):1277–1286. doi:10.1016/j.neuroimage.2012.03.068.

26. O’Reilly JX, Woolrich MW, Behrens TEJ, Smith SM, Johansen-Berg H. Tools of the trade: psychophysiological interactions and functional connectivity. Soc Cogn Affect Neurosci. 2012;7(5):604–9. doi:10.1093/scan/nss055.

27. Gitelman DR, Penny WD, Ashburner J, Friston KJ. Modeling regional and psychophysiologic interactions in fMRI: the importance of hemodynamic deconvolution. Neuroimage. 2003;19(1):200–207.

28. Field M, Cox WM. Attentional bias in addictive behaviors: a review of its development, causes, and consequences. Drug Alcohol Depend. 2008;97(1-2):1–20. doi:10.1016/j.drugalcdep.2008.03.030.

29. Stoeckel LE, Kim J, Weller RE, Cox JE, Cook EW, Horwitz B. Effective connectivity of a reward network in obese women. Brain Res Bull. 2009;79(6):388–95. doi:10.1016/j.brainresbull.2009.05.016.

30. Luo S, Monterosso JR, Sarpelleh K, Page KA. Differential effects of fructose versus glucose on brain and appetitive responses to food cues and decisions for food rewards. Proc Natl Acad Sci USA. 2015;112(20):6509–14. doi:10.1073/pnas.1503358112.

31. Richard JM, Castro DC, Difeliceantonio AG, Robinson MJF, Berridge KC. Mapping brain circuits of reward and motivation: in the footsteps of Ann Kelley. Neurosci Biobehav Rev. 2013;37(9 Pt A):1919–31. doi:10.1016/j.neubiorev.2012.12.008.

32. Robinson TE, Berridge KC. Review. The incentive sensitization theory of addiction: some current issues. Philos Trans R Soc L B Biol Sci. 2008;363(1507):3137–3146. doi:10.1098/rstb.2008.0093.

33. Aston-Jones G, Smith RJ, Sartor GC, et al. Lateral hypothalamic orexin/hypocretin neurons: A role in reward-seeking and addiction. Brain Res. 2010;1314:74-90. doi:10.1016/j.brainres.2009.09.106.

34. Ponz A, Khatami R, Poryazova R, et al. Abnormal activity in reward brain circuits in human narcolepsy with cataplexy. Ann Neurol. 2010;67(2):190–200. doi:10.1002/ana.21825.

35. Cocchi L, Zalesky A, Fornito A, Mattingley JB. Dynamic cooperation and competition between brain systems during cognitive control. Trends Cogn Sci. 2013;17(10):493–501. doi:10.1016/j.tics.2013.08.006.

36. Power JD, Petersen SE. Control-related systems in the human brain. Curr Opin Neurobiol. 2013;23(2):223–228. doi:10.1016/j.conb.2012.12.009.

37. Ridderinkhof KR, Ullsperger M, Crone EA, Nieuwenhuis S. The Role of the Medial Frontal Cortex in Cognitive Control. Science (80-). 2004;306(5695).

38. Delazer M, Högl B, Zamarian L, et al. Executive functions, information sampling, and decision making in narcolepsy with cataplexy. Neuropsychology. 2011;25(4):477–87. doi:10.1037/a0022357.

39. Zamarian L, Högl B, Delazer M, et al. Subjective deficits of attention, cognition and depression in patients with narcolepsy. Sleep Med. 2015;16(1):45–51. doi:10.1016/j.sleep.2014.07.025.

40. Renn RP, Cote KA. Performance monitoring following total sleep deprivation: Effects of task type and error rate. Int J Psychophysiol. 2013;88(1):64–73. doi:10.1016/j.ijpsycho.2013.01.013.

41. Drummond S, Brown GG. The Effects of Total Sleep Deprivation on Cerebral Responses to Cognitive Performance. Neuropsychopharmacology. 2001;25(5):S68–S73. doi:10.1016/S0893-133X(01)00325-6.

42. Thomas M, Sing H, Belenky G, et al. Neural basis of alertness and cognitive performance impairments during sleepiness. I. Effects of 24 h of sleep deprivation on waking human regional brain activity. J Sleep Res. 2000;9(4):335–52.

43. Papies EK, Stroebe W, Aarts H. Healthy cognition: processes of self-regulatory success in restrained eating. Pers Soc Psychol Bull. 2008;34(9):1290–300. doi:10.1177/0146167208320063.

44. Hollitt S, Kemps E, Tiggemann M, Smeets E, Mills JS. Components of attentional bias for food cues among restrained eaters. Appetite. 2010;54(2):309–13. doi:10.1016/j. appet.2009.12.005.

